# Archaeal lineages related to eukaryotes encode functional diterpenoid cyclases

**DOI:** 10.1101/2025.02.07.637177

**Authors:** H. Solomon McShea, Valerie De Anda, Jochen J. Brocks, Paula V. Welander, Brett J. Baker

## Abstract

The first eukaryotic cell originated through the union of an archaeon and a bacterium. Phylogenetic evidence suggests that the archaeal partner was a member of phylum Asgardarchaeota, likely within the Heimdallarchaeia clade. Although little is known about the molecular basis of eukaryogenesis, insight can be gained from studying the unique biochemistry of modern Asgardarchaeota. Here, we investigated the potential for Asgards to produce cyclic terpenoids. Phylogenomics coupled with structural prediction revealed that organisms from the clades Hodarchaeales and Kariarchaeaceae encode for diterpenoid cyclases. We tested the functionality of these enzymes *in vitro*, showing that they cyclize geranylgeranyl pyrophosphate to form bicyclic halimadienyl pyrophosphate. Halimadienyl nucleosides have previously been shown to mediate intracellular persistence of *Mycobacterium tuberculosis* in host endosomes, and may function similarly in Asgardarchaeota. The characterization of Asgard diterpenoid cyclases provides insight into the biochemistry of an elusive group of microbes that are pivotal in the evolution of complex cellular life.

## Main Text

The discovery and characterization of the Asgardarchaeota (written here as “Asgards,” also known as Promethearchaeati [1]) revolutionized our understanding of the origin of eukaryotes and eukaryotic cell biology [2,3]. Phylogenetic analysis has shown that Asgards are likely the closest archaeal relatives of eukaryotes [2,4], supporting the hypothesis that an Asgard ancestor was a partner in the endosymbiosis [5–7] from which eukaryotic life arose. Analysis of modern Asgard genomes suggests that these organisms have some archaeal-like physiological features such as isoprenoidal membranes [8], as well as features otherwise only known in eukaryotes such as membrane trafficking proteins and a dynamic cytoskeleton [9–11]. However, relatively little study has been devoted to Asgard proteins that are not ubiquitous in eukaryotes, despite the potential of these proteins to reveal unique Asgard biology, including processes that may have been important in eukaryogenesis.

In pursuit of such non-universal proteins, we identified 64 putative diterpenoid cyclase genes in archaeal genomes [15] and metagenome-assembled genomes (MAGs), including 31 from Asgards. All but three Asgard sequences are from Heimdallarchaeia, the clade thought to contain the Asgard ancestor of eukaryotes [4], and most are from hydrothermal vents (see Supplementary Methods and Table S1). Four are from organisms which have recently been isolated in co-culture [14]. All archaea, including Asgards, produce the diterpenoid cyclase substrate, geranylgeranyl pyrophosphate (GGPP), in the course of archaeal membrane lipid biosynthesis [12]. The polycyclic products of diterpenoid cyclases range in function from communication (e.g. gibberellins in plants and their symbionts) to conflict (e.g. platencin and terpentecin in soil bacteria) [13]. The relevance of these functions to Asgard symbiosis, especially the eukaryogenetic symbiosis, motivated us to determine the biosynthetic products of Asgard diterpenoid cyclases.

To determine the function of putative Asgard cyclases, we performed *in vitro* experiments with four Asgard proteins found in our initial search. As a control, we also tested the *Mycobacterium tuberculosis* H37Rv homolog, Rv3377c, which had been previously shown to produce halimadienyl pyrophosphate [16]. The Asgard cyclases include two homologs from *Candidatus* Hodarchaeales archaeon S146_22 (WAPZ01000470.1 and WAPZ01000551.1), one (OLS20947.1) from *Candidatus* Heimdallarchaeota archaeon LC2 (assigned to Kariarchaeaceae by the Genome Taxonomy Database), and one from *Candidatus* Lokiarchaeales archaeon CR 4 (OLS12628.1). These proteins have the acidic DxD motif [17] necessary to catalyze terpenoid cyclization (Fig. S1), a hydrogen bond donating residue at position H341 (corresponding to sequence position in Rv3377c), which positions the general acid for proton transfer, and a bulky aromatic residue at position W380 to protect the general acid when substrate is not bound.. We incubated the purified protein (Fig. S2) with geranylgeranyl pyrophosphate (GGPP; **1’**) and analyzed the reaction products by gas chromatography-mass spectrometry (GC-MS). We found that two of the Asgard cyclases (OLS20947.1 from *Candidatus* Heimdallarchaeota archaeon LC2 and WAPZ01000551.1 from *Candidatus* Hodarchaeales archaeon S146_22) convert GGPP to halimadienyl pyrophosphate (**2’**) (Fig. 1).

**Figure 1.**
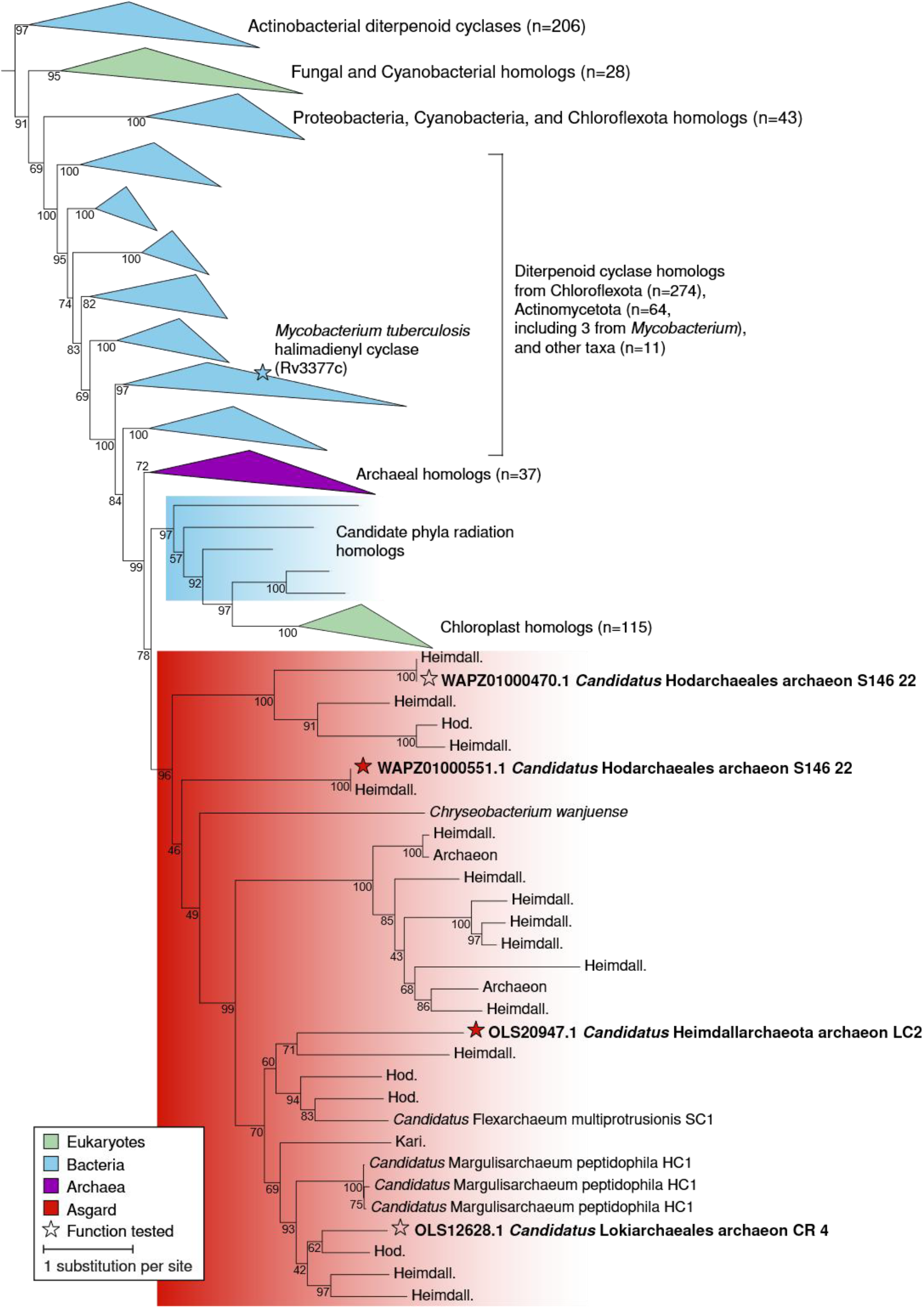
Asgard cyclase homologs are nested within the diterpenoid cyclases. The maximum likelihood phylogeny was rooted by treating all non-diterpenoid cyclases (e.g., sterol and hopanoid cyclases) as an outgroup (see [23]). Nodes are marked with ultrafast bootstrap support. Enzymes tested in this study are marked with stars, which are filled for those that were functional. Asgard homologs not tested are marked with abbreviations indicating Asgard phylum, where Heimdall. = Heimdallarchaeales, Hod. = Hodarchaeales, Kari = Kariarchaeaceae. Supplemental Data 1 contains this tree in Newick format.

Phylogenetic analysis (Fig. 2) revealed that Asgard sequences are monophyletic and sister diterpenoid cyclases from plant chloroplasts. The chloroplast clade has several bacterial sequences from the Candidate Phyla Radiation (CPR) on its stem. All sequences in the Asgard-dominated clade are from Heimdallarchaeia except for one from Lokiarchaeales and one from a bacterium. Sister to Asgard and plant sequences is a clade of archaeal sequences from Thermoplasmata, Theionarchaea, and Methanofastidiosia groups of superphylum Methanobacteriati (also known as Euryarchaeota), with lone sequences from the superphylum Thermoproteati (TACK) taxa Nitrososphaerota and Bathyarchaeia, two sequences from the Asgard phylum Thorarchaeota, and two from bacteria. Outside the archaeal-plant group, paraphyletic bacterial groups are dominated by Chloroflexota, particularly sequences from sponge symbiont MAGs, suggesting a possible bacterial source for sponge cyclic diterpenoids [18]. Among these groups are clusters of Actinomycetota sequences, including one which contains the characterized [16] halimadienyl cyclase Rv3377c from *M. tuberculosis*. The deepest-branching clade is large and composed primarily of bacterial sequences from Actinomycetota, including characterized proteins which synthesize precursors to gibberellins and terpenoid antibiotics [19,20]. This phylogeny shows that unlike the distantly related sterol and hopanoid cyclases, which are near-ubiquitous and perhaps ancestral in eukaryotes and bacteria respectively, diterpenoid cyclases are quite taxonomically restricted, but often widespread within those taxa.

**Figure 2.**
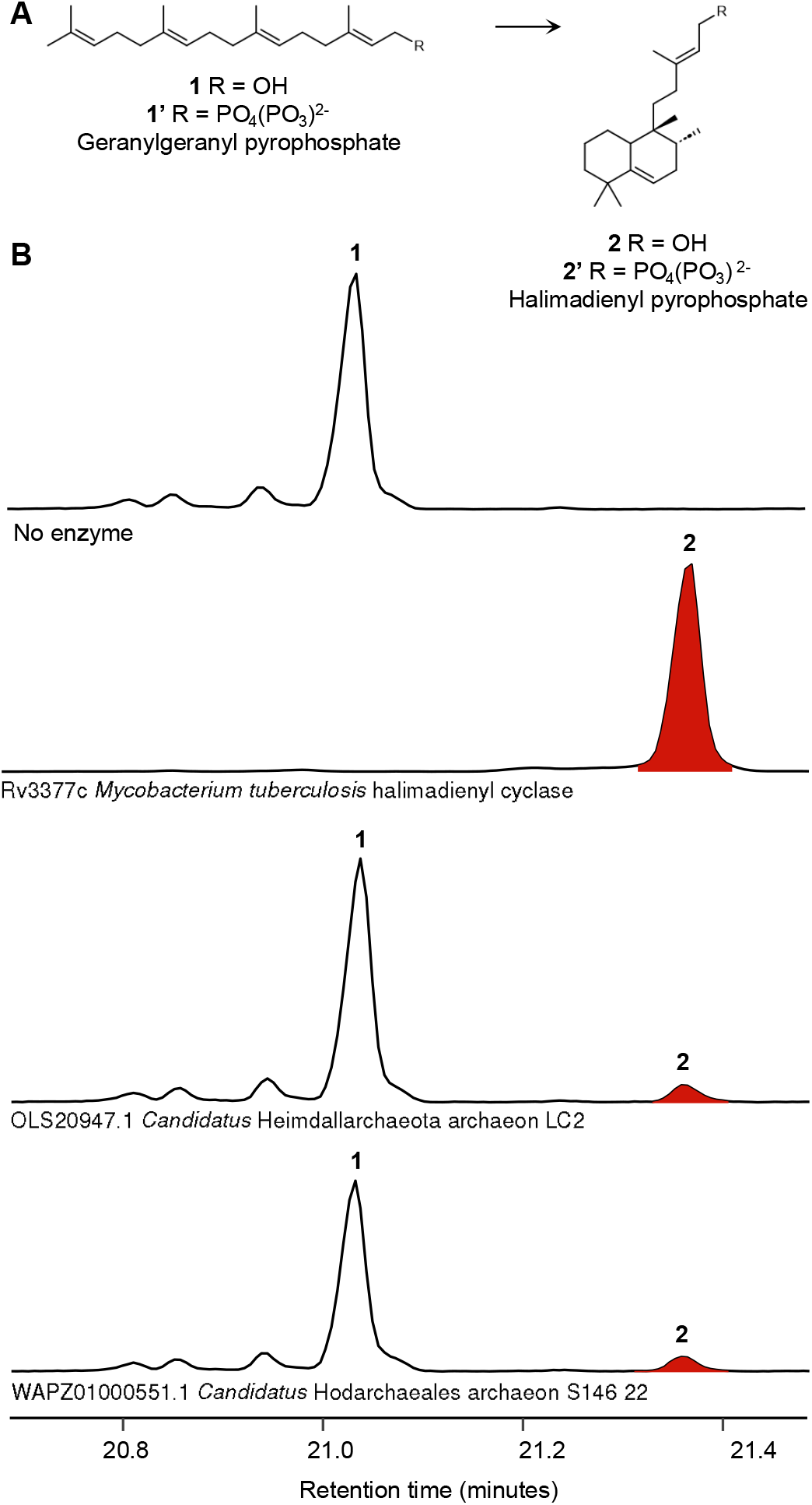
Asgard cyclases catalyze the cyclization of geranylgeranyl pyrophosphate (GGPP) to form halimadienyl pyrophosphate. (*A*) Cyclization reaction and dephosphorylation products, identified by comparison to published gas chromatography-mass spectrometry (GC-MS) spectra (Fig. S3). (*B*) Extracted ion chromatograms (*m/z* 119, 189, 191, 290, 362) of *in vitro* reactions performed with GGPP and cyclases. Reaction products were dephosphorylated by incubation with phosphatase, extracted with hexanes, and trimethylsilylated before GC-MS. The alcohols geranylgeraniol (**1**) and halimadienol (**2**) were only detected after dephosphorylation, suggesting that the cyclization product is halimadienyl pyrophosphate (**2’**).

Halimadienyl cyclases homologs are prevalent in Heimdallarchaeia (Hodarchaeales and Kariarchaeaceae) but rare in other Asgards. Heimdall cyclases could have been acquired vertically from a common ancestor with Methanobacteriati, with gene loss in other Asgard phyla, or via horizontal transfer on the Heimdall stem. The position of chloroplast cyclases sister to, rather than nested within, those from Asgards, suggests that these eukaryotes did not inherit their diterpenoid cyclases from Asgards. However, the position of plant sequences in the cyclase tree and that of eukaryotes within the Asgard genome tree may change as additional Asgards are sequenced. Regardless of evolutionary history, the enrichment of diterpenoid cyclases in Heimdallarchaeia may suggest a function for these enzymes that is unique to Heimdall biology, and that may have been shared by the Asgard ancestor of eukaryotes.

The biological role of halimadienyl compounds in Asgards is unknown. The only other microbe known to synthesize halimanes is *M. tuberculosis*, an intracellular bacterial pathogen of mammalian macrophages. *M. tuberculosis* constitutively secretes 1-halimadienyl-adenosine into the host endosome, where it acts as a Lewis base to neutralize endosomal pH and halt maturation of the endosome into a lysosome, facilitating survival and persistence of the bacterium [21]. In addition to the cyclase, 1-halimadienyl-adenosine biosynthesis requires an adenosyl transferase [22], which is also found in the genome of *Candidatus* Hodarchaeales S146_22. If this organism can synthesize the functional antacid, it could stabilize endosomal persistence of the Asgard in a host cell, or of symbionts acquired through an endomembrane trafficking system [3] in the Asgard cell. Testing these hypotheses will require characterizing the S146_22 adenosyl transferase homolog and its product, and ultimately (co-)culturing this Asgard to determine whether it can participate in endosymbiosis. Currently, endosymbiosis has not been observed in modern Asgards despite their ancestors’ likely role in the eukaryogenetic endosymbiosis. However, the biology of halimadienyl cyclases in Asgards, as well as that of other Asgard proteins not widespread in eukaryotes, holds great promise for understanding their unique biology and their potential role in cellular evolution.

## Acknowledgments

We thank Alysha Lee, Andy Garcia, Charles Hu, Maryam Khademian, Minh Tu, Tharika Liyanage, and Maggie Horst for helpful discussions, as well as Daniel Fernandez, Olivia Pattelli, and the Stanford Macromolecular Structure Knowledge Center (MSKC) for advice, enthusiasm, and facilities for protein purification. H.S.M. and P.V.W. were supported by NSF Grant EAR-1752564. H.S.M. was supported by the National Science Foundation (NSF) Graduate Research Fellowship, the Stanford Enhancing Diversity in Graduate Education (EDGE) Fellowship, the Stanford School of Sustainability McGee and Levorsen Graduate Research Grant, and the Stanford MSKC Training Program in Biophysical and Structural Analysis of Biological Macromolecules. Portions of the experiments were performed in the Stanford Geomicrobiology Shared Laboratories Core Facility (RRID:SCR_025000). Computational analyses were performed on the Sherlock high-performance computing cluster administered by the Stanford Research Computing Center. This work was also supported by the Moore-Simons Project on the Origin of the Eukaryotic Cell, Simons and Moore Foundation 73592LPI to B.J.B. (https://doi.org/10.46714/735925LPI).

## Author Contributions

H.S.M.: conceptualization, data curation, investigation, methodology, formal analysis, visualization, writing – original draft, writing – review & editing; V.D.A.: conceptualization, investigation, writing – review & editing; J.J.B.: conceptualization, writing – reviewing & editing; P.V.W.: conceptualization, supervision, writing – reviewing & editing; B.J.B.: conceptualization, supervision, writing – reviewing & editing.

