## Supplemental Information for "Archaeal lineages related to eukaryotes encode functional diterpenoid cyclases"

**Supplementary Methods, Figures, and Tables for**
Archaeal lineages related to eukaryotes encode functional diterpenoid cyclases.

H. Solomon McShea[1,2†], Valerie De Anda[3,4,5†], Jochen J. Brocks[6], Paula V. Welander[2*], and Brett J. Baker[3,4*]

[1]Department of Microbiology and Immunology, University of California, San Francisco, San Francisco, CA, USA
[2]Department of Earth System Science, Stanford University, Stanford, CA, USA
[3]Microbial Oceanography Unit, Department of Functional and Evolutionary Ecology, University of Vienna, Vienna, Austria
[4]Department of Marine Science, University of Texas Austin, Marine Science Institute, Port Aransas, TX, USA
[5]Department of Integrative Biology, University of Texas Austin, TX, USA
[6]Research School of Earth Sciences, Australian National University, Canberra, ACT, Australia
†Contributed equally

**Keywords:** Asgardarchaeota, diterpenoid cyclases, biosynthesis, eukaryogenesis.

### Supplementary Methods

**Terpenoid cyclase database curation.** We curated a Type II terpenoid cyclases database to analyze the phylogenetic position and evolutionary history of Asgard cyclases, following previous work on terpenoid cyclase phylogeny [1]. Briefly, we first used sequences from each known clade of Type II cyclases to query public databases: Joint Genome Institute Integrated Microbial Genomes & Microbiomes database (JGI IMG; https://img.jgi.doe.gov), the National Center for Biological Informatics (NCBI) nonredundant (nr), clustered nonredundant (clustered nr), sequence read archive (sra) and whole-genome shotgun contigs (wgs) databases (https://www.ncbi.nlm.nih.gov/genbank), and the European Molecular Biology Laboratory – European Bioinformatics Institute (EMBL-EBI) MGnify database (https://www.ebi.ac.uk/metagenomics). JGI IMG and NCBI were queried via BLAST [2], with an expect threshold of 0.05 and a word size of 5 for NCBI, and an e-value cutoff of 1e-5 for JGI. MGnify was queried via phmmer [3] with an e-value cutoff of 1e-5. We also downloaded all hits for Pfams PF13243 (squalene-hopene cyclase C-terminal domain) and PF13249 (squalene-hopene cyclase N-terminal domain) from EBI (https://www.ebi.ac.uk/interpro/entry/pfam/#table). These searches returned 44,209 unique sequences, which, after a length filter of 350 (the length of the smallest known Type II cyclase [4]) to 900 (the length of the largest known Type II domain-bearing cyclase [5]) and subsetting by CD-hit (command cd-hit -i in.fasta -o out.fasta -c 0.90 -n 5 -M 6000 -d 0 -T 8) [6], yielded a database of 13,680 sequences. Finally, after initial phylogenies were estimated and the split between diterpenoid and triterpenoid cyclases was apparent, we created a custom hidden Markov models (HMM) using domain-wise (β and γ) alignments of the diterpenoid cyclase group. The diterpenoid β domain model was very similar to the triterpenoid β domain model as measured by overlapping HMMER [7] hits on the whole database. In contrast, the diterpenoid γ domain model had no reciprocal hits with the triterpenoid γ domain model. These diterpenoid cyclase-specific models were used to search our databases again, returning 6 additional diterpenoid cyclase homologs from Asgards. Later BLAST searches on the NCBI nr and wgs databases returned an additional 21 Asgard homologs.

**Phylogenetic estimation and analysis of terpenoid cyclases.** Cyclase sequences were aligned using the MAFFT linsi algorithm [8] implemented in Magus [9]. The alignment was trimmed using the ClipKIT [10] kpic-gappy algorithm (command clipkit in.aln -m kpic-gappy -o out.aln). The trimmed alignment was then used to create an HMM, with which we extracted the shared cyclase domain from full-length sequences. Cyclase domains were then aligned and trimmed, and phylogenies were estimated with IQ-TREE v2.2.2.6 or v3.0.1 [11–14]. The best-fit model of amino acid sequence evolution, EX_EHO + R10, was determined using multiple ModelFinder [15] runs to completely explore model space, and branch support was calculated with ≤ 10,000 ultrafast bootstrap approximations [16]. Tree visualization was performed in Dendroscope 3.8.10 [17] and iTOL 6.8.2 [18].

**Heterologous gene expression and protein purification.** Four Asgard cyclase gene sequences, along with positive control Rv3377c from *M. tuberculosis*, were codon-optimized for expression in *Escherichia coli* and synthesized by Twist Bioscience (South San Francisco, CA) in a pET-28a(+) plasmid (Table S2). Plasmids sequences were confirmed by Plasmidsaurus (Eugene, OR) and plasmids were transformed by heat shock into competent NiCo21(DE3) cells (New England Biolabs) for expression, along with pACYC for overexpression of the *E. coli* GroEL/ES chaperone system and Trigger Factor (Addgene #83923) (Table S3). As previously reported for Rv3377c [19], Asgard cyclases were only functional when expressed alongside these folding chaperones. Media was supplemented with kanamycin (15 µg/mL) and chloramphenicol (20 µg/mL) to maintain selection for both plasmids.

Expression strains were cultured in biological triplicate in 1 L terrific broth (TB) in a 2 L flask. Cultures were grown at 37 °C while shaking at 225 rpm to an $OD_{600}$ of ~0.6, cooled on ice, induced with 500 µM isopropyl β-D-1-thiogalactopyranoside (IPTG), then grown an additional ~20 h at 16 °C shaking at 225 rpm. Cells were harvested by centrifugation at 10,000 × g. Pellets were stored at −20 °C or immediately resuspended in 30 mL lysis buffer (50 mM Tris, pH 7.5, 2 mM $MgCl_2$, 0.5 mM tris(2-carboxyethyl)phosphine, 10% glycerol, 0.1% Triton X-100, 0.3 M NaCl, 0.2 mg/mL lysozyme, 0.3 mL Xpert Protease Inhibitor Cocktail; modified from [20]) for sonication. Disruption by sonication was achieved with a 6.4 mm microtip at 50% amplitude on a 30 s on, 30 s off cycle for 5 minutes of on time. Lysates were cleared by centrifugation at 15,000 × g.

Protein was purified by immobilized metal affinity chromatography with a 5 mL HisTrap FF Nickel column (Cytiva Life Sciences) on an Akta Pure (General Electric) fast protein liquid chromatograph (FPLC). After equilibration with loading buffer (50mM Tris, pH 7.5; 0.1 M NaCl), the sample was loaded at 0.5 mL/min, and elution buffer (50mM Tris, pH 7.5; 0.1 M NaCl, 0.5 M imidazole) concentration was increased to 100% at 4%/min. Protein was further purified by size-exclusion chromatography, and purity was assessed via SDS-PAGE.

***In vitro* enzymatic activity assay and workup.** Enzymatic reactions were prepared by combining 20 µg enzyme and 50 µM geranylgeranyl pyrophosphate (GGPP) (Sigma-Aldrich) in 2.5 mL of buffer (50 mM Tris, pH 7.5, 0.1 mM MgCl2, 0.1% Tween-80, after [19]). The reactions were incubated at 30 °C for 18 h. Reaction products were dephosphorylated with the addition of 50 µL Quick CIP (and 250 µL rCutsmart buffer, both from New England Biolabs) and incubation at 37 °C for 20 m. The reactions were then extracted three times with an equal volume of hexanes, with centrifugation at 2,800 × g before each extraction to facilitate separation of polar and nonpolar phases. Extracts were dried under a gentle stream of $N_2$ gas and subsequently derivatized to trimethylsilyl ethers by the addition of 50 µL pyridine and 50 µL *N,O*-Bis(trimethylsilyl) trifluoroacetamide for 1 h at 70 °C prior to GC-MS analysis.

**Product determination by GC-MS.** Organic extracts were separated on an Agilent 7890B Series gas chromatograph (GC) with helium as the carrier gas at a constant flow of 1.1 ml/min. The GC program ran as follows: 50 °C for 3 minutes, ramp 14 °C /min to 300 °C, hold 3 min, ramp 10 °C /min to 330 °C, and hold for 5 min (modified from [21]). Separation was achieved using two tandem DB17-HT columns (30 m × 0.25 mm i.d. × 0.15 µm film thickness). 2 µL of sample were injected into a Gerstel-programmable temperature vaporization (PTV) injector operated in splitless mode at 250 °C. The GC was coupled to a 5977A Series mass selective detector (MSD) with the source at 320 °C and operated in electron ionization (EI) mode scanning from 90 to 600 Da in 0.3 s. Compounds were identified by comparing mass spectra to previously published spectra [22–25].

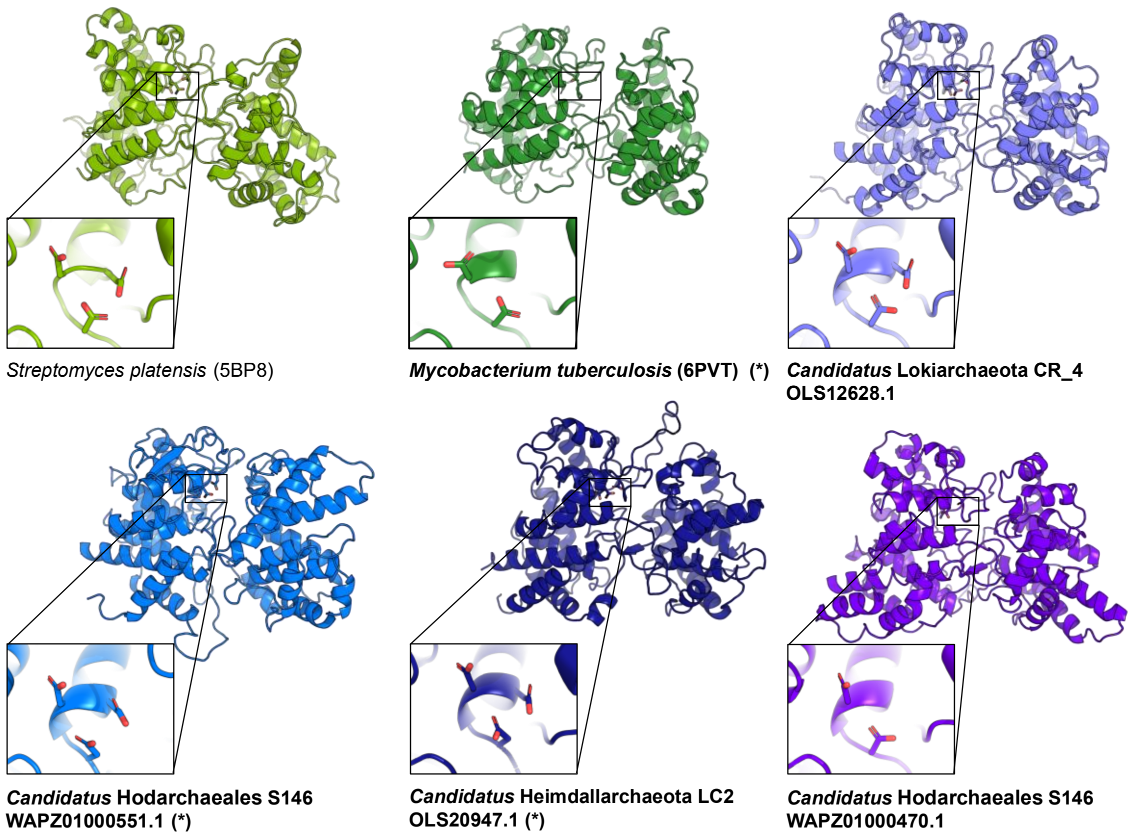
*Streptomyces platensis* (5BP8)
***Mycobacterium tuberculosis*** **(6PVT) (*)**
***Candidatus*** **Lokiarchaeota CR_4**
**OLS12628.1**
***Candidatus*** **Hodarchaeales S146**
**WAPZ01000551.1 (*)**
***Candidatus*** **Heimdallarchaeota LC2**
**OLS20947.1 (*)**
***Candidatus*** **Hodarchaeales S146**
**WAPZ01000470.1**

| | |
|---|---|
| **Rv3377c Mycobacterium tuberculosis H37Rv (*)** | GVGWTGNSTLE**DCD**TTSVAYDVLSKFG |
| **WAPZ01000470.1 Candidatus Hodarchaeales archaeon S146 22** | HLSYASTLRIY**DLDD**TLATYIGLKLLG |
| **WAPZ01000551.1 Candidatus Hodarchaeales archaeon S146 22 (*)** | GIGISPFF-IP**DAD**AIAMALQDALLLN |
| **OLS12628.1 MBAA01000202.1 Candidatus Lokiarchaeales archaeon CR 4** | GLAFGKFFSIA**DSDD**TSLGLYLLEKYG |
| **OLS20947.1 MDVR01000113.1 Candidatus Heimdallarchaeota archaeon LC2 (*)** | GLAFNGNFPFT**DSDD**TSLGLYLMDLTN |
| BEU34980.1 AP029002.1 Candidatus Margulisarchaeum peptidophila HC1 | GIGWSDNFPVS**DADD**TSVGLYLLHDYR |
| BEU35449.1 AP029002.1 Candidatus Margulisarchaeum peptidophila HC1 | GIGWSDNFPVS**DADD**TSVGLYLLHDYR |
| BEU40373.1 AP029002.1 Candidatus Margulisarchaeum peptidophila HC1 | GIGWSDNFPVS**DADD**TSVGLYLLHDYR |
| BFF49627.1 AP029247.1 Candidatus Flexarchaeum multiprotrusionis SC1 | GIGFSKHFPLP**DFDD**TIVALNLLQSNK |
| DAHWUS010000547.1 Candidatus Heimdallarchaeota archaeon clean1540 | GVGHSSCFPMT**DADD**TALSLLLLNKFY |
| JAJRQD010000059.1 Archaeon Dive100 bin10.34 | GISFGYSFPIL**DADD**TSVALVVLKKLG |
| JAJRTV010000293.1 Archaeon Dive96 bin10.124 | GISLSEYFDIC**DSDD**TIVLMKFLENED |
| JAJRWK010000064.1 Archaeon Dive96 bin2.176 | GVPFGESFPIP**DADD**TAVAIVVLANLG |
| JALTSK010000185.1 Candidatus Heimdallarchaeota archaeon S140 metabat2 scaf2bin.067 | GLSFGKAFKVP**DADD**TSVALVDLHYMG |
| JALUHM010000709.1 Candidatus Heimdallarchaeota archaeon S144 maxbin2 scaf2bin.078 | GIGISPFF-IP**DAD**AIAMALQDALLLN |
| JALUIC010001379.1 Candidatus Heimdallarchaeota archaeon S144 maxbin2 scaf2bin.193 | WISAASTYPVI**DLDD**TAVAHLTLGRLG |
| JALUOM010000943.1 Candidatus Heimdallarchaeota archaeon S146 maxbin2 scaf2bin.134 | HLSYASTLRIY**DLDD**TLATYIGLKLLG |
| JALUPF010000472.1 Candidatus Heimdallarchaeota archaeon S146 maxbin2 scaf2bin.220 VB | GVPFGESFPIS**DADD**TAMAIVVLASLG |
| JALUPQ010000268.1 Candidatus Heimdallarchaeota archaeon S146 maxbin2 scaf2bin.295 | LLGFSSSFNVP**DLDD**TLMVHLVLRRLG |
| JALUTL010000122.1 Candidatus Heimdallarchaeota archaeon S146 metabat2 scaf2bin.271 | GLSFGKGFMNP**DAD**NTAVALIDLIYMG |
| JASBSB010000004.1 Candidatus Heimdallarchaeota archaeon BWSM 123 | GLSLGKSFQIP**DSDD**TSIALLLLDKYS |
| JBJKSQ010000558.1 Candidatus Hodarchaeales archaeon Bin 86 | GIGFGEHFTIQ**DVDD**TALALKLLNDNN |
| JBPMHF010000063.1 Candidatus Hodarchaeales archaeon 3336-2 bin.17 | WLGFSDSYDVP**DLDD**TLMVHLILRRCG |
| JBPMSN010000066.1 Candidatus Kariarchaeaceae archaeon 357-6-R1-1-358 bin.53 | GLGFAKDF-YS**DSD**TTSLAILLIHTNG |
| MDH5403948.1 JAOUKW010000326.1 Candidatus Heimdallarchaeota archaeon 8 1 Jan SF Bin58 | GIPFGKYFPIP**DADD**TAVALYVLHHYG |
| MFV2016789.1 JBGMRU010001563.1 Candidatus Heimdallarchaeota archaeon N04 concoct2 bin.40 | GVGHSQYFPMT**DADD**TSVSLYLFEKYK |
| MFW9878233.1 JBHLRM010000858.1 Candidatus Thorarchaeota archaeon TM1M.329 | GISMSKLYYYE**DLDD**TATVFYLLNQTN |
| MFX0095459.1 JBHLKR010001053.1 Candidatus Hodarchaeaota archaeon DZ1M.13 | GVSFSEHFPVP**DLDD**SVLSLYLLTKYE |
| MGD2072663.1 JASFTW010000107.1 Candidatus Thorarchaeota archaeon D1103U20metaG sub Bin 339 | GVGWSTELPVT**DADD**TALACRALCAMD |
| MHA2364396.1 JBSWQY010000106.1 Candidatus Hodarchaeales archaeon M3-38 Bin 173 | GVGYSKHISLT**DADD**TVLGLHLLEQHG |
| WAKA01000042.1 Candidatus Heimdallarchaeota archaeon S012 26 esom | GLSFGKAFPVP**DADD**TAVAIVDMHYMG |
| WAMJ01000055.1 Candidatus Heimdallarchaeota archaeon S139 21 | GISFGYSFPVL**DADD**TAIALLVKNYLG |

**Supplementary Figure 1:** Asgard diterpenoid cyclases have canonical terpenoid cyclase macromolecular structure (top) and active site motif (bottom; this motif is DxD[D] in diterpenoid cyclases, DxDD in squalene-hopene cyclases, and xxDC in sterol cyclases). Cyclases tested *in vitro* are marked in bold in both panels; those that synthesized halimadienyl pyrophosphate are marked with an asterisk (*). Structures were predicted using Alphafold2 [26], alignment was estimated with MAFFT [8].

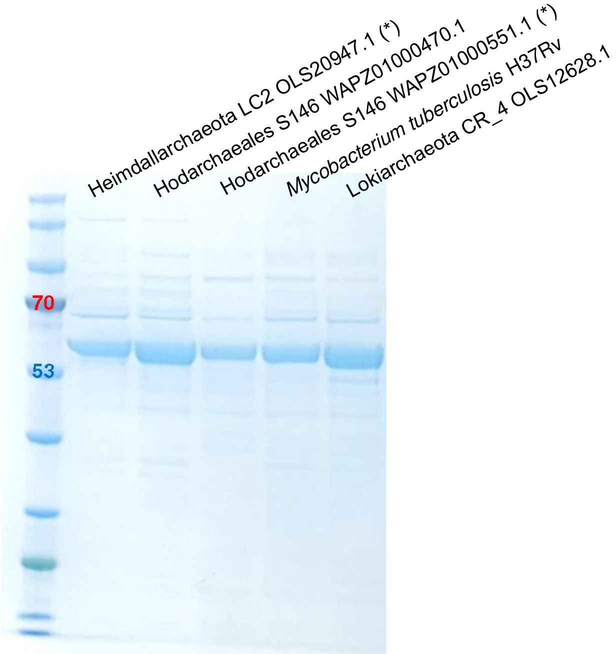


**Supplementary Figure 2:** Asgard cyclases and positive control from *M. tuberculosis* were partially purified by affinity chromatography.

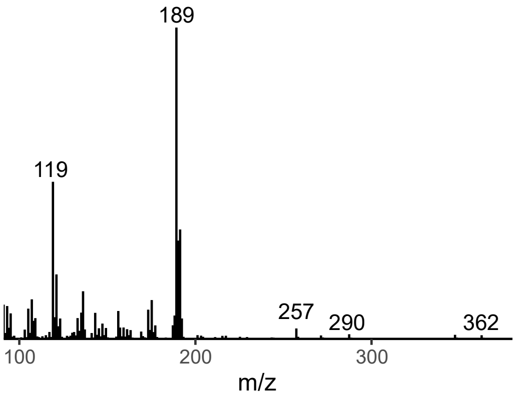


**Supplementary Figure 3:** Mass spectrum of peak at 21.35 minutes, identified by comparison to published spectra as the trimethylsilyl derivative of 2, halimadienol.

**Supplementary Table 1 (attached .xlsx):** Asgard cyclases identified and/or tested for GGPP cyclization in this study.

**Supplementary Table 2:** Plasmids used in this study.

| Plasmid | Description | Source |
|---|---|---|
| pET-28a(+) | pET derivative with ColE1/pMB1/pBR322/pUC ori, Kan$^R$, N-terminal 6x His tag. All cyclases were cloned into this plasmid. | Twist Bioscience |
| pACYC-GroEL/ES-TF (pPVW7383) | pACYC-Duet1 derivative with p15A ori, Cam$^R$. Addgene #83923. Expresses GroEL/ES and trigger factor. | [27] |

**Supplementary Table 3:** Strains constructed in this study.

| Expression strain | Plasmids |
|---|---|
| HM138 | pACYC (pPVW7383), pET-Mycobacterium_tuberculosis_Rv3377c (pHM115) |
| HM141 | pACYC (pPVW7383), pET-OLS20947.1_Heimdallarchaeota_LC2 (pHM118) |
| HM144 | pACYC (pPVW7383), pET- WAPZ01000551.1_Hodarchaeales_S146 (pHM134) |
| HM145 | pACYC (pPVW7383), pET-WAPZ01000470.1_Hodarchaeales_S146 (pHM135) |
| HM147 | pACYC (pPVW7383), pET- OLS12628.1_Lokiarchaeales_CR_4 (pHM119) |
